# Animal soundscapes reveal key markers of Amazon forest degradation from fire and logging

**DOI:** 10.1101/2021.02.11.430853

**Authors:** Danielle I. Rappaport, Anshuman Swain, William F. Fagan, Ralph Dubayah, Douglas C. Morton

**Affiliations:** Department of Geographical Sciences, University of Maryland, College Park MD 20742; Department of Biological Sciences, University of Maryland, College Park MD 20742; Biospheric Sciences Laboratory, NASA Goddard Space Flight Center Greenbelt, MD 20771

**Keywords:** Bioacoustics, ecosystem services, land use, habitat loss, forest disturbance, frontier forests

## Abstract

Safeguarding tropical forest biodiversity requires solutions for monitoring ecosystem composition over time. In the Amazon, logging and fire reduce forest carbon stocks and alter tree species diversity, but the long-term consequences for wildlife remain unclear, especially for lesser-known taxa. Here, we combined data from multi-day acoustic surveys, airborne lidar, and satellite timeseries covering logged and burned forests (n=39) in the southern Brazilian Amazon to identify acoustic markers of degradation. Our findings contradict theoretical expectations from the Acoustic Niche Hypothesis that animal communities in more degraded habitats occupy fewer ‘acoustic niches.’ Instead, we found that habitat structure (e.g., aboveground biomass) was not a consistent proxy for biodiversity based on divergent patterns of acoustic space occupancy (ASO) in logged and burned forests. Full 24-hr soundscapes highlighted a stark and sustained reorganization in community structure after multiple fires; animal communication networks were quieter, more homogenous, and less acoustically integrated in forests burned multiple times than in logged or once-burned forests. These findings demonstrate strong biodiversity co-benefits from protecting Amazon forests from recurrent fire activity. By contrast, soundscape changes after logging were subtle and more consistent with community recovery than reassembly. In both logged and burned forests, insects were the dominant acoustic markers of degradation, particularly during midday and nighttime hours that are not typically sampled by traditional field surveys of biodiversity. The acoustic fingerprints of degradation history were conserved across replicate recording locations at each site, indicating that soundscapes offer a robust, taxonomically inclusive solution for tracking changes in community composition over time.

**Significance Statement:** Fire and logging reduce the carbon stored in Amazon forests, but little is known about how human degradation alters animal communities. We recorded thousands of hours of ecosystem sounds to investigate animal community assembly and the associations between biodiversity and biomass following Amazon forest degradation over time. 24-hr patterns of acoustic activity differed between logged and burned forests, and we observed large and sustained breakpoints in community structure after multiple burns. Soundscape differences among degraded forests were clearest during insect-dominated hours rarely sampled in field studies of biodiversity. These findings demonstrate that acoustic monitoring holds promise for routine biodiversity accounting, even by non-experts, to capture a holistic measure of animal communities in degraded tropical forests and benchmark change over time.

## Introduction

Biological diversity is disappearing rapidly in response to human activity, especially in tropical forests, home to well over half of Earth’s terrestrial species (1). Global concern over greenhouse gas emissions from tropical forests (2) has led to international efforts to Reduce Emissions from Deforestation and Forest Degradation (REDD+) (3). Retention of diverse ecosystems supports climate change mitigation and adaptation (4); and yet, carbon-focused conservation may not result in a commensurate win for tropical forest biodiversity (5). Longstanding data gaps on species distributions and uncertainty regarding the direct and indirect impacts of human activity on biodiversity complicate efforts to quantify the interplay between carbon and biodiversity (6).

Across the tropics, the Brazilian Amazon has the highest rates of deforestation (7), and forest degradation from fire and logging may double biodiversity loss from deforestation alone (8). However, the long-term impacts of human activity on Amazon biodiversity remain highly uncertain, due, in part, to the spatial heterogeneity among degraded forests from differences in the timing, frequency, extent, and severity of disturbances (9).Time-varying heterogeneity in the biodiversity of degraded forests may also explain some of the apparent contradictions in previous studies of degradation impacts on birds, the most well studied Amazonian taxa. Nectarivorous birds, for example, increase in abundance immediately after logging but ultimately decline, whereas insectivorous birds show immediate sensitivity to changes in habitat from logging and continue to decline in abundance over time (10). Time dependence has also complicated efforts to measure the effects of degradation on insects, a problem confounded by limited research (11).

Addressing the tropical biodiversity crisis therefore requires an efficient, distributed, long-term monitoring system to assess species abundance and diversity (12). Traditional, ground-based biodiversity inventories are logistically prohibitive to conduct at scale, and limited taxonomic expertise perpetuates large data discrepancies for lesser-known taxa, such as insects, which constitute the bulk of tropical biodiversity (6). Advances in the emerging discipline of acoustic remote sensing, or *ecoacoustics,* may permit large-scale biodiversity monitoring for multiple taxa, including unidentifiable species, based on the aggregate sound signature of the animal community, or soundscape (13–15). Since multiple sites can be recorded simultaneously over time, sound surveys reduce the effort and cost associated with routine monitoring and facilitate standardized assessments of community variation and ecosystem recovery. Most previous efforts to utilize acoustic data for biodiversity monitoring have focused on detecting known vocalizations associated with individual species (16, 17), but there is increasing interest in evaluating the entire collection of signals in a given soundscape to derive measures of ecosystem intactness that include all sound-generating taxa without definitive species identification (14, 15, 18, 19).

The Acoustic Niche Hypothesis (ANH) (20) is a core premise of ecoacoustics and the prevailing organizing principle for assessing species richness (15), beta diversity (21), and human impacts (22, 23) from soundscape data. The ANH posits that more biodiverse habitats exhibit finer niche partitioning of available transmission space among species, as described by frequency and time of day, and thus, greater acoustic space occupancy (ASO). The corollary is that more degraded habitats support less acoustic infilling due to more vacant ‘acoustic niches’ from local species extirpations (24). Ecoacoustic approaches have great potential to extend monitoring capabilities in the hyperdiverse tropics, where competition for acoustic space is strongest (25). Still, large uncertainties remain as to whether acoustic space infilling can be used as a robust proxy for ecosystem intactness to monitor landscapes altered by human activity (26).

Here, we test the ANH across logged and burned Amazon forests to identify acoustic markers of forest degradation (Fig. 1). We collected coincident high-density airborne lidar data and multi-day acoustic recordings (1192.5 hours) during September-October 2016 in 39 forests with different times-since-logging (4-23 years) and histories of fire activity (1-5 fires), stratified based on a 33-year time series of annual Landsat imagery (9). We used space-for-time substitution and two complementary analytic approaches to characterize threshold effects and time dependence for the composition and connectivity of animal acoustic networks along gradients of degradation history (see Materials & Methods). First, we calculated ASO for each site at hourly and one-minute time steps to test the ANH and to quantify the magnitude, variability, and persistence of shifts in animal community structure following forest degradation. Second, we developed a novel network-based approach to capture additional complexity from the soundscape data to track the composition and co-occurrences of ‘acoustic pseudo-taxa’ (defined as clusters of organisms that coordinate acoustic activity by time and frequency) along degradation and recovery pathways. Our findings demonstrate that soundscapes encode digital markers of degradation and reveal distinct patterns of community assembly following logging and fire, paving the way for more widespread use of ecoacoustics to benchmark and monitor changes in animal community composition in human-altered tropical forest landscapes.

**Figure 1.**
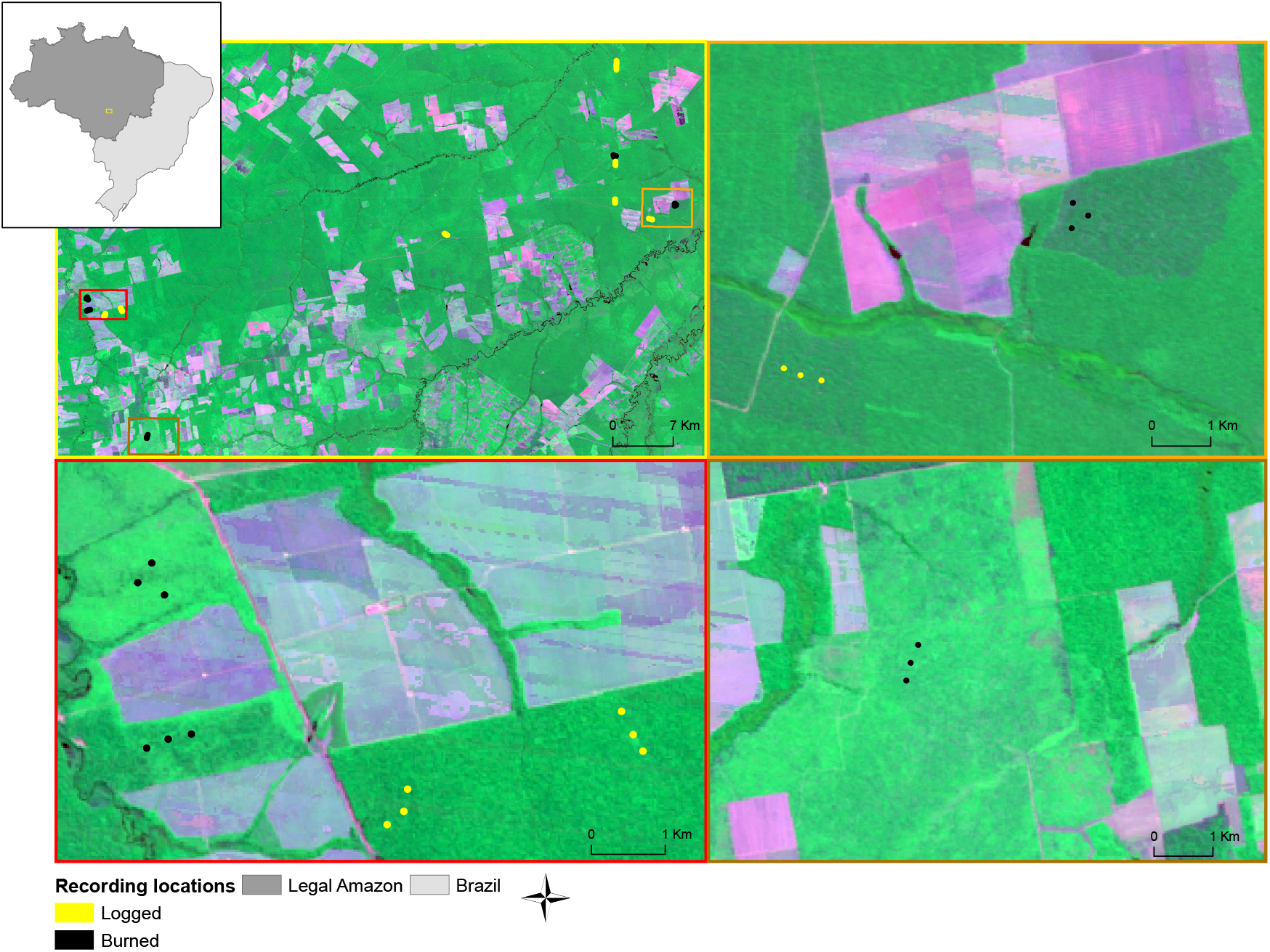
Acoustic recording sites in logged and burned forests (n=39) were distributed across 9,400 km^2^ in northern Mato Grosso, Brazil (upper left). Colored boxes identify subsets of the study domain to illustrate how the triplicate sampling design captures the heterogeneity in habitat structure and acoustic community composition in degraded forests. False-color composites of Landsat imagery (2014, 543-RGB) in each panel show deforested areas in magenta and gradients of forest cover in shades of green.

## Results

Soundscape data from degraded Amazon forests did not support the ANH (Fig. 2). Instead, acoustic analyses revealed contrasting impacts of fire and logging on community structure. After fire, daily ASO increased linearly with estimated aboveground biomass, whereas daily ASO in logged forests did not increase with biomass (Fig. 2). Importantly, ASO-degradation relationships varied with time of day (Fig. 2; Fig. S1), highlighting the value of full-day measurements from autonomous recording devices for detailed investigations of the ANH and the dominant indicators of degradation.

**Figure 2.**
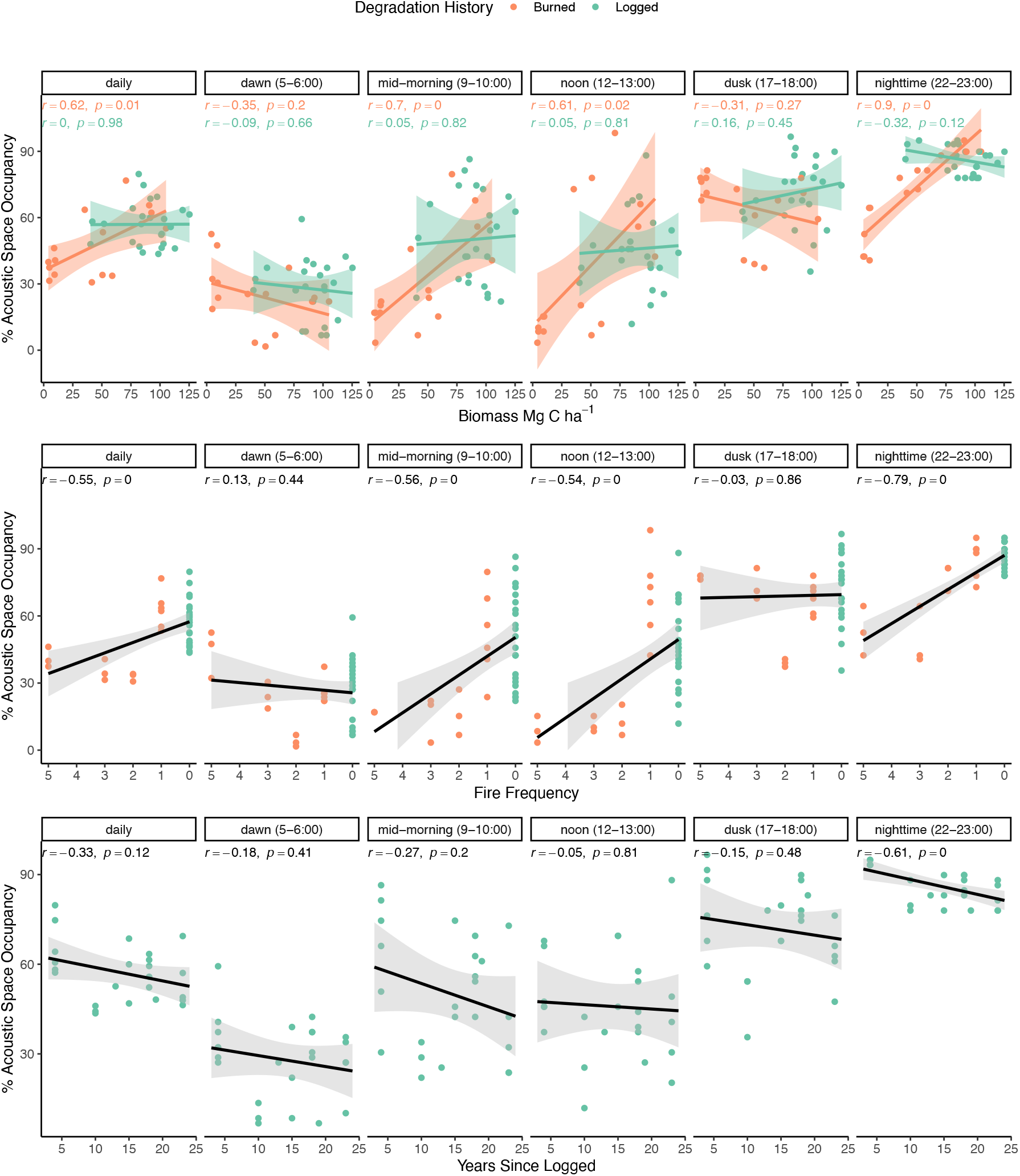
Patterns of soundscape infilling did not conform to expectations from the Acoustic Niche Hypothesis that animal communities in more intact habitats occupy more of the 24-hour soundscape, with intactness measured as either aboveground biomass (top), fire frequency (middle), or years-since-logging (bottom). Individual panels summarize Acoustic Space Occupancy (ASO) over specific time intervals of known biological activity for birds and insects to pinpoint likely taxonomic contributions to daily trends. Fig S1 provides the full 24-hour profiles for logged and burned forests, and Fig. S2 illustrates ASO relationships aggregated at 1-minute resolution.

Insects were the most obvious acoustic markers of changing community composition in burned forests. ASO during insect-dominated periods of the day (e.g., mid-morning, noon, nighttime) strongly differentiated burned forests as a function of both biomass (max |r| = 0.9 at 22:00-23:00) and fire frequency (max |r| = 0.82 at 20:00-21:00), and these time periods governed the overall daily trend (Fig. 2). Notably, ASO relationships with biomass and fire frequency were weakest during the 5:00-6:00 dawn chorus typical of bird surveys (p > 0.05; Fig. 2). In logged forests, the only time interval that exhibited a moderately strong relationship with logging age (max |r| = 0.61 at 22:00-23:00) showed an unexpected decline in ASO with increasing time since logging. ASO and biomass in logged forests were not correlated for any time period. Relationships between ASO and degradation history were consistent when aggregating acoustic niches at both hourly and minute time scales (Fig. S2).

The 24-hr profile of ASO and peaks of acoustic activity varied markedly among burned forests as a function of fire frequency and aboveground biomass, and less so among logged forests, despite a large range (50%) in aboveground biomass (Fig. 3, Fig. S3). The timing and magnitude of the dominant acoustic activity peaks were similar among logged forests (Fig. 3), which varied broadly in terms of time since logging (4-23 years) and potential impacts of logging infrastructure (e.g., skid trails, tree-fall gaps) (Fig. 1). Both logged and once-burned forests exhibited greater daily variability and overall use of acoustic space (ASO) than recurrently burned forests (Fig. 3, Fig. S3). Acoustic communities in recurrently burned forests occupied the least amount of frequency space during all time periods except dusk, which was the most heavily occupied time window for all degradation classes (17:00-18:00). Dawn hours had the lowest ASO across all 24-hour soundscapes except in the most heavily degraded forests burned 3 or more times.

**Figure 3.**
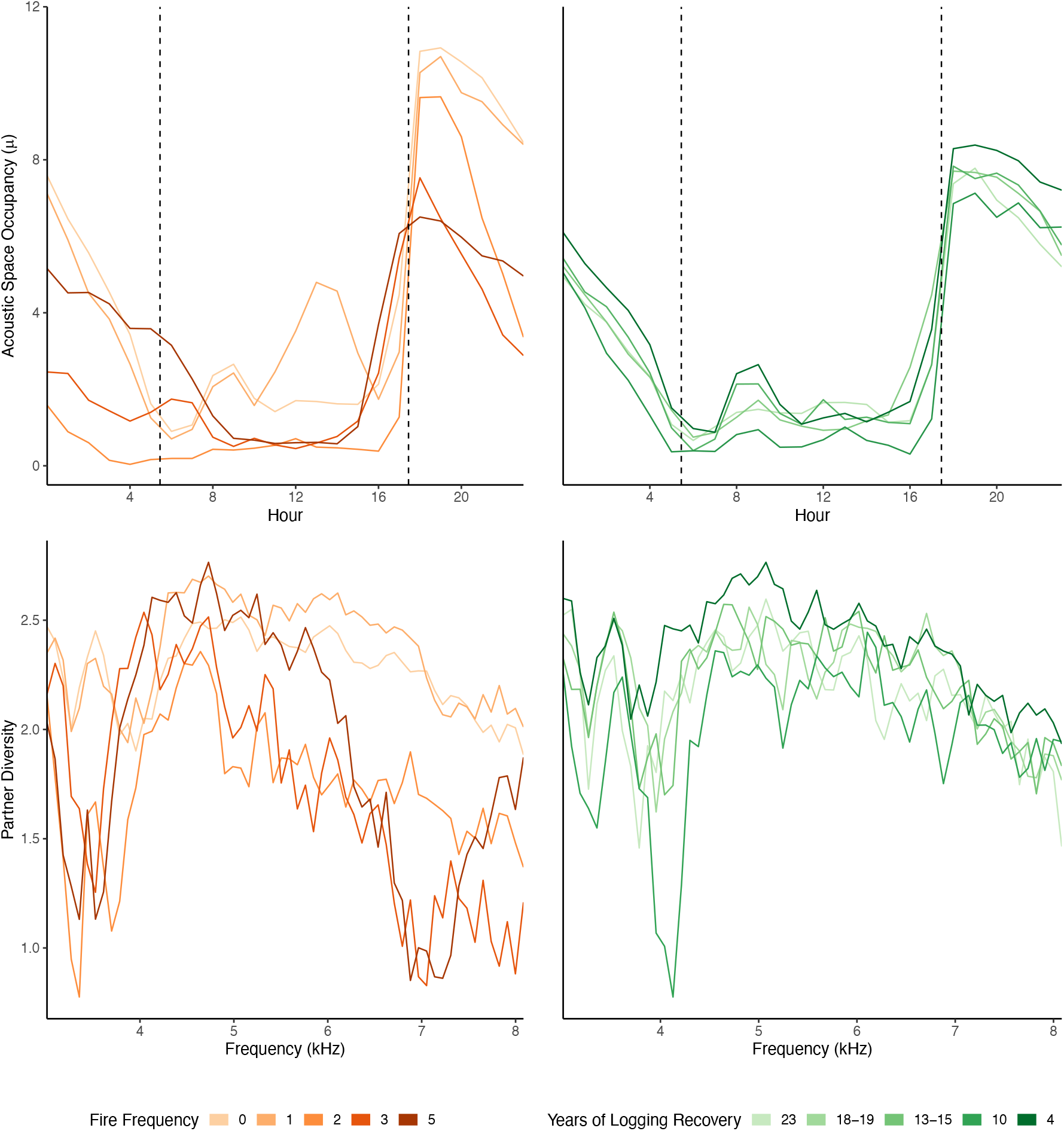
Soundscape data reveal distinct shifts in acoustic activity following recurrent burning (orange) and more subtle differences in logged forests (green). Top, mean daily profiles of Acoustic Space Occupancy (ASO). Bottom, *partner diversity,* a network-based analysis of the sound frequency bands that appear and disappear across gradients in fire frequency and time since logging. Colored lines represent the average response per degradation stratum, and dashed vertical lines indicate sunrise and sunset. Results for each recording site are shown in Figs S3 (ASO and *partner diversity* of sound frequency bands) and Fig. S4 *(partner diversity* of sound hours).

We identified a breakpoint in community composition between forests burned once and forests burned two or more times. This threshold effect from recurrent fire activity manifested as a sustained reduction in ASO from late morning through late afternoon (10:00-15:00; Fig. 3, Fig. S3). Differences across levels of initial fire severity were also most evident during this time interval, reflecting localized impacts of burn intensity on acoustic communities, even at short length scales (300 m) within the same fire event (Fig. 4). In contrast to the non-linear shifts observed midday, ASO declined linearly with increasing fire frequency after dusk (Fig. 3).

**Figure 4.**
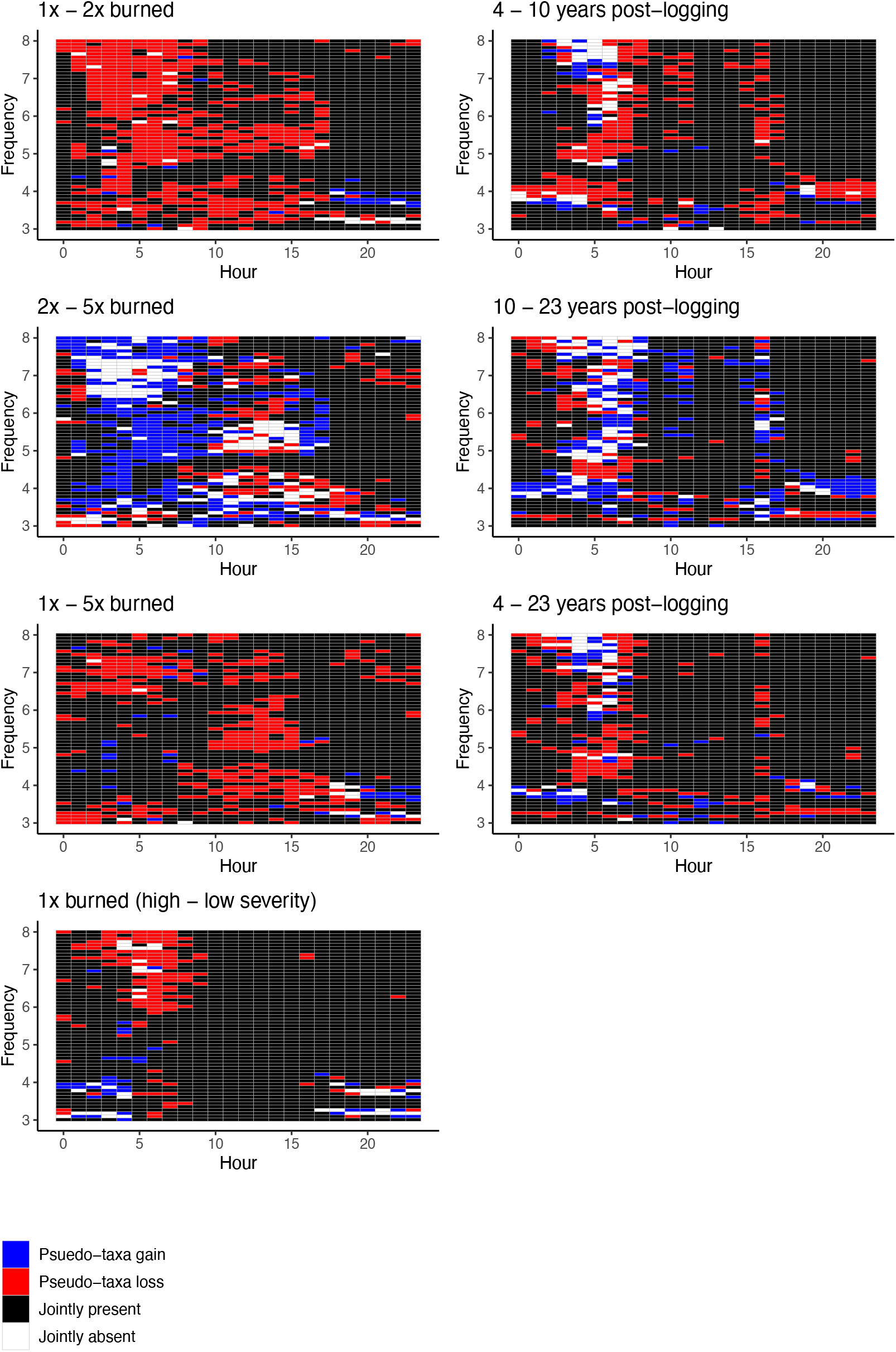
Pairwise differences in matrices of Acoustic Space Occupancy (ASO) identify coordinated losses and gains of pseudo-taxa as a function of degradation frequency, timing, and severity. Acoustic pseudo-taxa are gained (blue) or lost (red) in the transition between the first set of forests and the second set of forests in each labeled pair.

Network analyses indicated that differences in ASO reflected differences in community composition following logging and fire. Most frequencies, except the mid-range channels (~ 3.7–5.2 kHz), were detected during a greater diversity of hours in logged and once-burned forests than in recurrently burned forests, and thus had higher *partner diversity* (see Materials & Methods) (Fig. 3, Fig. S3). Importantly, the frequency bands, or acoustic pseudo-taxa, that best differentiated burned forests as a function of fire frequency (~3.5 kHz, 6.5-7.5 kHz) were not the same as those that best differentiated logged forests as a function of time since logging (~4.2 kHz, Fig. 3, Fig. S3). After logging, several acoustic pseudo-taxa appeared and disappeared with recovery time, but most were conserved across logged forest sites with 4-23 years of regrowth (Fig. 4). For the majority of frequency bands, their greatest *partner diversity* was observed following 4 years of regeneration, followed by a sharp but short-term decline between 4 and 10 years. The dynamic range of network attributes among logged classes was much more limited than among burned classes.

Overall, fire resulted in more empty acoustic niches across the 24-hour soundscape than logging, and recurrent burns led to a major restructuring of the animal community. Sets of acoustic pseudo-taxa were lost and not regained between 1-2 fires and 1-5 fires (Fig. 4). In all transitions, losses and gains were clustered in time-frequency space. For example, a large cluster of losses during the transition from 1 to 5 fires was detected midday (10:00-15:00), the same time period shown in Fig. 2 that clearly differentiated forests by fire frequency. Animal communication networks also became more acoustically homogenous with increasing fire recurrence. *Alatalo interaction evenness,* a measure of signal homogeneity across time-frequency space, increased linearly with increasing fire frequency, while variance declined linearly among replicates within each class with increasing fire frequency (Fig. 5). The evenness of the communication network further explained variation in successional recovery after fire: *Alatalo evenness* was consistently lower in younger once-burned stands than older once-burned stands, reflecting an increased dominance of fewer sets of sounds.

**Figure 5.**
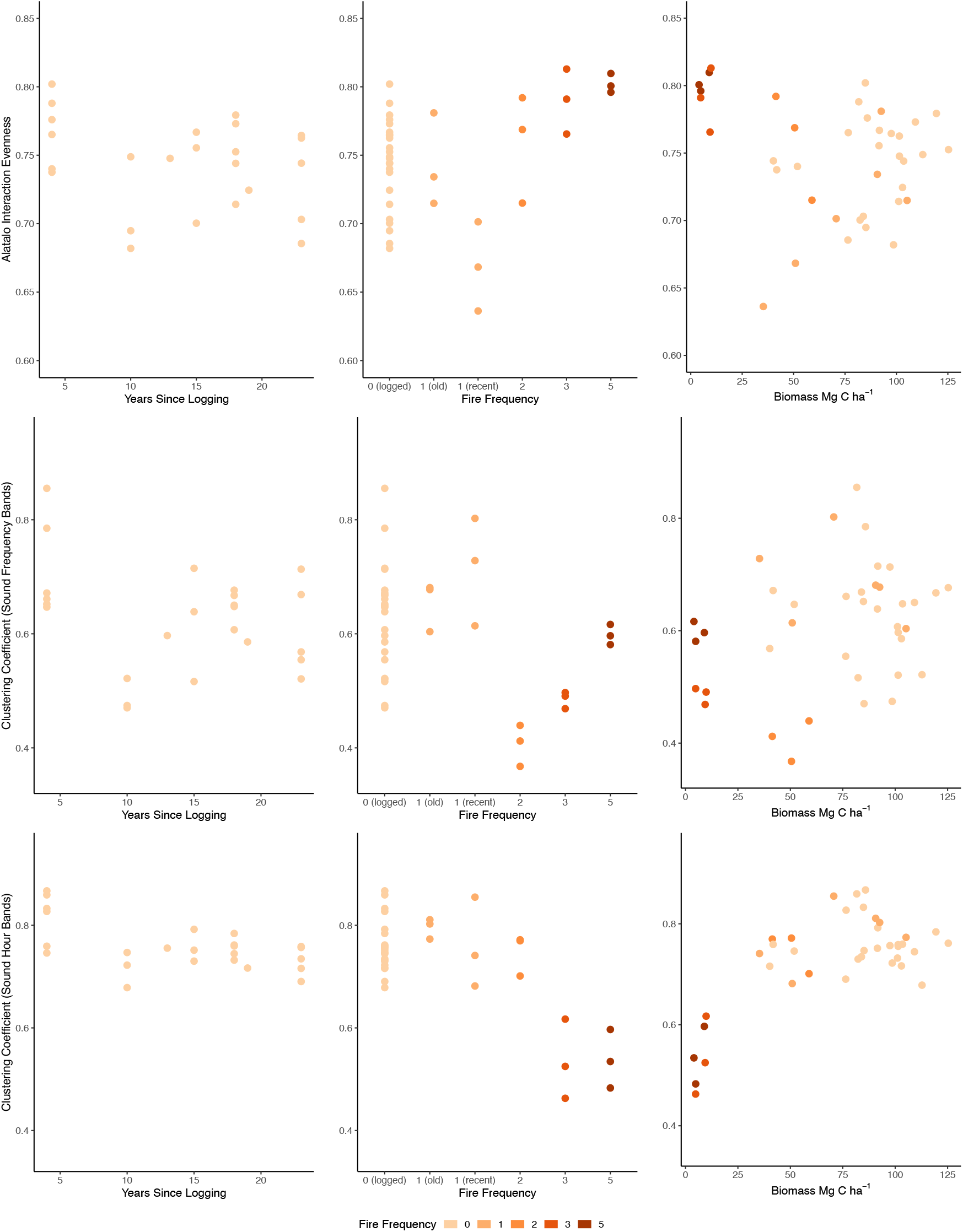
Forest degradation alters animal communication networks in frontier Amazon forests, as measured by *Alatalo interaction evenness* (top), which quantifies the homogenization and dominance of signals across the animal communication network, defined by time and frequency (top), and the *Clustering coefficient,* the tendency for acoustic pseudo-taxa to form coordinated clusters over frequency (middle) and time (bottom).

Closely clustered sound groups, or ‘acoustic guilds,’ from local-scale network metrics such as the *clustering coefficient* (which measures transitivity in co-occurrences of acoustic pseudo-taxa over frequency or time), offer a more synthetic understanding of the component processes that drive system-level patterns, including ASO and evenness. The large drop in the *clustering coefficient* of frequency bands between 1 and 2 fires indicated a disintegration of time-synchronized groups of spectrally similar sounds (Fig. 5). The subsequent increase in the clustering tendency of acoustic pseudo-taxa between 2 and 5 fires was of a fundamentally different character, as new acoustic guilds appeared at frequencies that had limited overlap with the acoustic guilds lost between 1 and 2 fires (Figs. 4–5).

By contrast, animal communication networks after logging suggested community-level recovery rather than the reorganization observed due to fires. Several acoustic guilds appeared and disappeared with time since logging (e.g., 4 kHz), but most were conserved across logged forests of different successional stages (Figs. 3–4). Many time-frequency niches that went silent between 4 and 10 years after logging were re-occupied between 10 and 23 years (Fig. 4). We observed a short-term decline in *Alatalo interaction evenness* and the *clustering coefficient* of sound hours and sound frequency bands between 4 and 10 years after logging (Fig. 5). However, this effect appeared transient, with soundscapes of logged forests far less variable than fire-affected soundscapes, based on the similarity in ASO and network patterns, and the transition matrix of losses and gains of acoustic pseudo-taxa (Figs. 3–5).

## Discussion

Acoustic soundscapes capture the full 24-hr profile of animal activity in forest ecosystems to provide a taxonomically holistic assessment of community changes and reassembly following human degradation. Results from this study expand upon previous field studies restricted to specific taxa over limited survey periods and habitat ranges (27–29). Soundscape data highlighted distinct trajectories of community structure after logging and fire, which did not correlate with aboveground biomass. Differences in acoustic community composition between burned and logged forests underscore the need to track long-term changes in tropical forest biodiversity to complement carbon monitoring systems such as REDD+. In particular, we found major restructuring of the animal community after two or more fires. Soundscape markers of forest degradation were strongest and most consistent during midday and midnight insect choruses, time periods rarely sampled in field-based inventories of birds, the most common study taxa (30). By revealing the hidden structure of acoustic assemblages, the network-based methods in this study offer a new window into the nature of animal communities in frontier Amazon forests. Network analyses of ecoacoustic remote sensing data offer a scalable approach to track changes in community assembly over space and time.

Patterns of ASO following logging and fire did not vary with aboveground biomass, a common metric of tropical forest intactness (31), and part of the theoretical basis for the ANH. Instead, we found that animal communities in less intact forests did not contain more soundscape gaps (i.e., empty acoustic niches). One possible explanation is that the long-term evolutionary mechanisms that favor acoustic niche partitioning may lose relevance during transient periods as animal communities respond to or reorganize following disturbances (32). Further work that directly compares acoustic community structure before and after forest degradation may help to resolve this inconsistency. Surveys in forests without a history of logging or fire may also reveal additional insights about the ANH and baseline patterns of ASO, although intact forests are rare in frontier Amazon landscapes. Longer-term surveys of ASO may also help interrogate the ANH across broader seasonal scales, such as during the rainy season with greater acoustic sensitivity to amphibians. Independent of the ANH, consistent acoustic fingerprints among replicate sites and daily surveys suggest that network-based analyses of soundscapes may help differentiate among transient typologies of forest succession. These results corroborate previous reports based on acoustic studies of birds that soundscapes function as memory banks of Amazon forest modification (33), but this study goes further to demonstrate that soundscapes may be most valuable when collected as 24-hr records that capture more acoustically active taxa and their network-level interactions.

Insects (e.g., orthopterans, hemipterans), not birds, were the strongest acoustic markers of threshold changes in degraded forest communities, highlighting the value of nighttime surveys for a more complete picture of biodiversity in human-modified forests. By contrast, soundscape measures from bird-dominated dawn and dusk intervals did not differentiate degraded frontier forests, possibly because bird choruses decline rapidly in response to even moderate perturbations to forest cover (22). In Borneo, degraded forest soundscapes were also found to be noisier and more homogenous at night, which was attributed to an influx of generalist insect species following logging (21). The major contribution of insects to the soundscape record in this study, and their sensitivity to habitat structure, may further explain why support for the ANH diverged from theoretical predictions and was inconsistent across degraded Amazon forests.

The mechanisms for partitioning acoustic space to maximize signal transmission vary among the major sound producers. For example, cricket species finely partition acoustic space by frequency, while tropical cicadas emit broadband signals that occupy wide frequency bands, such that neighboring species stratify acoustic space vertically, from the ground to canopy top, to avoid signal overlap (34, 35). Consequently, cicadas have evolved to occupy extremely narrow thermal niches that make them immediately responsive to changes in forest structure from human disturbance. Importantly, the timing of the noisy cicada chorus, which is governed by temperature and other conditions, affects the periods of acoustic activity by other species (36). Given that fire has more substantial impacts on forest structure than logging (8), commensurate changes in forest microclimate may therefore contribute to restructuring of the entire 24-hr animal communication network in burned forests (Fig. 5).

We found that animal communication networks in Amazon forests became quieter, less connected, and more homogenous after degradation from multiple fire events. Acoustic network analyses revealed a coordinated loss and gain of sounds (i.e., acoustic guilds) that represent distinct trajectories of biotic reorganization following repeated burns in Amazon forests. Evidence for successional divergence following repeated burning is consistent with experimental manipulation (37) and traditional species inventories (38). Because Amazonian forests are not adapted to fire, species that do survive initial or repeated burning can influence recovery pathways. For example, bird assemblages in Amazon forests burned multiple times are almost completely distinct from nearby unburned forests (39). In this study, the linear change in *Alatalo interaction evenness* after each recurrent fire event indicated biotic homogenization across all acoustic pseudo-taxa. Further, the *clustering coefficient* may be a particularly useful marker of degradation because it is sensitive to biological mechanisms that occupy broad swaths of acoustic space (e.g., cicadas, multi-species insect choruses). These results underscore the potential for further adaptation of network measures (40) to monitor community composition and soundscape changes over time.

Acoustic survey data in logged Amazon forests suggest a pattern of community recovery, rather than reassembly. Logging reduced the grouping tendency of acoustic pseudo-taxa and the *partner diversity* per sound frequency band, but only in the short term. Smaller effect sizes in ASO or network metrics from logging than fire in this study are consistent with the selective nature of logging in Amazon forests and more limited changes in forest structure (41) and thermal environment (42) than forest degradation from fire. The effect of logging on tropical forest biota is governed less by time-since-recovery and more by logging intensity (43), which may be less variable in the Amazon than other tropical forests. Overall, our findings are consistent with evidence for retention of guild structure following logging based on species-level responses from field studies (27).

Ecoacoustic surveys provide four main advantages compared to traditional field surveys for routine biodiversity monitoring in tropical forests. First, network-based measures of acoustic community composition provide a tractable solution for analyzing large volumes of recording data with an intermediate level of complexity between simple diversity indices and individual indicator species (44, 45). Second, autonomous survey equipment facilitates soundscape measurements over the full 24-hour cycle to capture lesser-known taxa (e.g., insects, amphibians) and time periods that best differentiate among degraded forests (e.g., midday, midnight). Third, sound sensors generate an objective, repeatable, and digital record that simplifies long-term monitoring and change detection over seasonal or interannual time scales. Finally, soundscape surveys are scalable because monitoring can be implemented by experts and non-experts alike, and acoustic data can be analyzed repeatedly ex-post to derive complementary information at different levels of granularity (e.g., community turnover, species populations) (17). Given these practical advantages, acoustic monitoring represents a promising low-cost solution for benchmarking current ecosystem conditions and monitoring biotic composition through time to augment space-based monitoring of tropical forest cover and habitat structure.

Acoustic measurements strongly differentiate between burned and logged Amazon forests, and provide clear evidence for breakpoints in community composition following recurrent burning. Space-based measures of forest habitat may not capture important differences in tropical forest biodiversity. Acoustic monitoring is therefore a complementary and efficient means to track changes over time from human activity at the tropical forest frontier. Given the potential biodiversity benefits of protecting burned Amazon forests from additional fires, this partnership between satellite and acoustic monitoring may support more robust biodiversity safeguards for REDD+ and related tropical forest conservation efforts. The integrity of Amazon forest ecosystems hinges upon policy and management that effectively prevents fire spread, particularly in frontier forest regions.

## Materials and Methods

### Remote sensing data

We selected 39 degraded Amazon forest study sites using a 33-year timeseries of annual Landsat imagery (1984-2017) to track the history of understory fire and selective logging in the municipalities of Nova Ubiritã and Feliz Natal in the Brazilian state of Mato Grosso (Fig. 1) (9). Logged sites (n=24) were sampled between 4 and 23 years post-disturbance, and once-burned sites were sampled between 5 (n=3) and 17 (n=3) years post-disturbance. Recurrently burned sites had 2 (n=3), 3 (n=3) or 5 (n=3) fires prior to airborne lidar and acoustic survey acquisitions. High-density airborne lidar data were acquired between 2013 and 2016 (≥ 14 returns per m^2^) (data available from: www.paisagenslidar.cnptia.embrapa.br/webgis/), and a regional model calibrated with forest inventory data in frontier forests was used to convert top of canopy height estimates to aboveground biomass (41) for each study site.

### Acoustic surveys and data processing

Circular forest sites (50 m radius) with uniform degradation history (forest patches ≥ 300 m radius) were sited at least 300 m from other recording sites or the forest edge. Most degradation strata contained three spatially proximate sites to capture the characteristic heterogeneity in forest structure at short length scales from logging infrastructure and differences in fire severity even within a single burn scar (9). Passive ARBIMON recorders (16) were installed at the center of each site to record one minute of sound every five minutes over 2-8 days during September-October 2016, totaling over 1100 hours of data. Sound detections were aggregated into three-dimensional soundscapes (site-level daily summaries of acoustic space use by the animal community) by binning signal detections into ‘acoustic niches’ defined by frequency (bin size: 83.13 Hz) and time. Two sets of 3D soundscape matrices were generated, based first on the native minute resolution and second on hourly resolution (x = time, y = frequency, z = presence/absence of biotic activity), representing the synoptic structure of the acoustic community for each site and daily survey. Analyses were constrained to frequencies between 3-8 kHz, which represent the greatest spectral overlap among bird, insect, and amphibian signals (15) and the spectral range with lowest likelihood for detection failure from acoustic attenuation in Amazon forests (19). We opted not to statistically correct for detection bias to avoid the large data loss from conforming to the assumptions of existing soundscape detection models that require all recordings with rain detections to be omitted from analysis (19). Instead, we restricted our analysis to the spectral domain least vulnerable to the potential confounding influence of forest structure, and we used an amplitude-filtering threshold of 0.2 to account for abiotic noise like rain when evaluating the occupancy status of each time-frequency channel (z-axis = binary presence/absence of biotic activity). Based on the two scales of aggregation, we evaluated 16992 channels (x-axis = 288 1-minute bins, y-axis = 59 frequency bins) and 1416 channels (x-axis = 24 1-hour bins, y-axis = 59 frequency bins). Acoustic space occupancy (ASO) was calculated based on the proportion of occupied time-frequency niches for each time step (mean ASO for soundscapes at native resolution and cumulative ASO for hourly soundscapes). A correlation analysis was used to evaluate habitat and ASO relationships at the scale of the day and constituent time intervals.

### Network analyses

Network construction and analyses were limited to frequencies between 3-8 kHz. We constructed weighted bipartite networks with sound frequency bins and sound hour bins as two classes of nodes. Frequency bins consisted of 60 nodes (3-8 kHz with bin size of 83.13 Hz) and time bins had 24 nodes (each depicting a 1-hour time interval during the day). Links between the two classes of nodes depicted presence of a given sound frequency bin during a noted sound hour bin and were weighted according to average number of observations for the link per day. Two levels of network metrics were constructed for each of the 39 sites in the dataset: the global-level analyses summarized the overall time-frequency topology of acoustic communication networks at a given site *(Alatalo interaction evenness* (46)), and the local-level analyses (node/class) unmixed ASO into the constituent elements that drive overall differences in network structure and connectivity *(Clustering coefficient* (47), *Partner diversity* (48)). The network analyses were performed with the aid of the following packages in R: ‘igraph’ (49), ‘vegan’ (50) and ‘bipartite’ (48).

*Alatalo interaction evenness* (AIE) measures heterogeneity in interactions across the network. Here, we focus on the frequency bins as total *n* entries, with pk are proportions of interactions of bin k, and calculate the metric as:

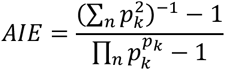

*Clustering coefficient* (CC) measures transitivity in co-occurrences/connection among the nodes of the same class. It can be calculated for the whole network, the class level as well as the node level. Here, we evaluated the *clustering coefficient* separately for sound frequency bins and sound hour bins by averaging the *clustering coefficients* of all nodes in a given class (i.e., rows and columns separately). It refers to the degree to which adjacent nodes in a graph tend to cluster together. In other words, if a frequency bin is present in two or more time bins, how many other frequency bins also share the same and vice-versa. It is based on the idea of triplets (47), which consist of three nodes that are joined either via two (open triplet) or three (closed triplet) undirected ties. The *clustering coefficient* is defined as:

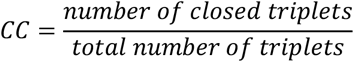

*Partner diversity* (PD) is the Shannon diversity of the number of interactions for a given node:

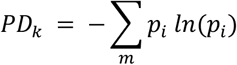

where, *PD_kl_* is the *partner diversity* of node *k* of a given class, which has *m* weighted connections from the other class of nodes, each of which has a proportion of interaction *p_i_* for a node *i* from the other bipartite node class. This value of PD can be calculated as prescribed for a node and then averaged for a given class, weighted by their marginal totals. PD can also be calculated for the entire network by weighted average of all the nodes’ PD values. *Partner diversity* of sound hours is consistent with ASO (Fig. S4).

## Acknowledgments

We gratefully acknowledge the support from the NASA Earth and Space Science Fellowship program and NASA’s Biological Diversity and Ecological Forecasting Program to D. I. Rappaport, and NASA’s Carbon Monitoring System to D.C. Morton. We also recognize support from the National Science Foundation through the Doctoral Dissertation Research Improvement Grant to D. I. Rappaport, and through award DGE-1632976 to A. Swain. Lidar data were acquired by Sustainable Landscapes Brazil with support from the US State Department, US Agency for International Development, and USDA-Forest Service. Marcos Longo, Maiza Dos-Santos and Hyeungu Choi provided assistance with lidar data processing, and Eveline Salvático provided field support for soundscape surveys in this study. We would also like to acknowledge Michael Keller and Matthew Hansen for their support and advising.

## References

1. T. A. Gardner, et al., Prospects for tropical forest biodiversity in a human-modified world. Ecology Letters 12, 561–582 (2009).

2. G. R. Van der Werf, et al., CO2 emissions from forest loss. Nature Geoscience 2, 737–738 (2009).

3. United Nations Framework Convention on Climate Change 2021 REDD+ Platform (January 30, 2021).

4. N. Seddon, B. Turner, P. Berry, A. Chausson, C. A. J. Girardin, Grounding nature-based climate solutions in sound biodiversity science. Nature Clim Change 9, 84–87 (2019).

5. J. Ferreira, et al., Carbon-focused conservation may fail to protect the most biodiverse tropical forests. Nature Climate Change 8, 744 (2018).

6. C. Meyer, H. Kreft, R. Guralnick, W. Jetz, Global priorities for an effective information basis of biodiversity distributions. Nature Communications 6, 8221 (2015).

7. P. G. Curtis, C. M. Slay, N. L. Harris, A. Tyukavina, M. C. Hansen, Classifying drivers of global forest loss. Science 361, 1108–1111 (2018).

8. J. Barlow, et al., Anthropogenic disturbance in tropical forests can double biodiversity loss from deforestation. Nature 535, 144–147 (2016).

9. D. I. Rappaport, et al., Quantifying long-term changes in carbon stocks and forest structure from Amazon forest degradation. Environ. Res. Lett. 13, 065013 (2018).

10. Z. Burivalova, et al., Avian responses to selective logging shaped by species traits and logging practices. Proceedings of the Royal Society of London B: Biological Sciences 282, 20150164 (2015).

11. F. França, et al., Do space-for-time assessments underestimate the impacts of logging on tropical biodiversity? An Amazonian case study using dung beetles. J Appl Ecol 53, 1098–1105 (2016).

12. M. Dornelas, et al., Towards a macroscope: Leveraging technology to transform the breadth, scale and resolution of macroecological data. Global Ecol Biogeogr 28, 1937–1948 (2019).

13. R. Gibb, E. Browning, P. Glover-Kapfer, K. E. Jones, Emerging opportunities and challenges for passive acoustics in ecological assessment and monitoring. Methods in Ecology and Evolution (2018).

14. S. S. Sethi, et al., Characterizing soundscapes across diverse ecosystems using a universal acoustic feature set. Proc Natl Acad Sci USA 117, 17049–17055 (2020).

15. T. M. Aide, A. Hernández-Serna, M. Campos-Cerqueira, O. Acevedo-Charry, J. L. Deichmann, Species Richness (of Insects) Drives the Use of Acoustic Space in the Tropics. Remote Sensing 9, 1096 (2017).

16. T. M. Aide, et al., Real-time bioacoustics monitoring and automated species identification. PeerJ 1, e103 (2013).

17. J. LeBien, et al., A pipeline for identification of bird and frog species in tropical soundscape recordings using a convolutional neural network. Ecological Informatics 59, 101113 (2020).

18. J. Sueur, A. Farina, A. Gasc, N. Pieretti, S. Pavoine, Acoustic indices for biodiversity assessment and landscape investigation. Acta Acustica united with Acustica 100, 772–781 (2014).

19. D. I. Rappaport, J. A. Royle, D. C. Morton, Acoustic space occupancy: Combining ecoacoustics and lidar to model biodiversity variation and detection bias across heterogeneous landscapes. Ecological Indicators 113, 106172 (2020).

20. B. Krause, Bioacoustics, habitat ambience in ecological balance. Whole Earth Review 57, 14–18 (1987).

21. Z. Burivalova, et al., Using soundscapes to investigate homogenization of tropical forest diversity in selectively logged forests. J Appl Ecol 56, 2493–2504 (2019).

22. Z. Burivalova, et al., Using soundscapes to detect variable degrees of human influence on tropical forests in Papua New Guinea. Conservation Biology 32, 205–215 (2018).

23. J. L. Deichmann, A. Hernández-Serna, J. A. Delgado C., M. Campos-Cerqueira, T. M. Aide, Soundscape analysis and acoustic monitoring document impacts of natural gas exploration on biodiversity in a tropical forest. Ecological Indicators 74, 39–48 (2017).

24. S. L. Dumyahn, B. C. Pijanowski, Soundscape conservation. Landscape Ecology 26, 1327–1344 (2011).

25. R. Planqué, H. Slabbekoorn, Spectral Overlap in Songs and Temporal Avoidance in a Peruvian Bird Assemblage. Ethology 114, 262–271 (2008).

26. A. Eldridge, et al., Sounding out ecoacoustic metrics: avian species richness is predicted by acoustic indices in temperate but not tropical habitats. Ecological Indicators (2018).

27. J. Barlow, C. A. Peres, L. M. P. Henriques, P. C. Stouffer, J. M. Wunderle, The responses of understorey birds to forest fragmentation, logging and wildfires: an Amazonian synthesis. Biological Conservation 128, 182–192 (2006).

28. R. B. de Andrade, et al., Tropical forest fires and biodiversity: dung beetle community and biomass responses in a northern Brazilian Amazon forest. J Insect Conserv 18, 1097–1104 (2014).

29. N. G. Moura, et al., Idiosyncratic responses of Amazonian birds to primary forest disturbance. Oecologia, 1–14 (2015).

30. J. Troudet, P. Grandcolas, A. Blin, R. Vignes-Lebbe, F. Legendre, Taxonomic bias in biodiversity data and societal preferences. Scientific Reports 7, 9132 (2017).

31. S. Frolking, et al., Forest disturbance and recovery: A general review in the context of spaceborne remote sensing of impacts on aboveground biomass and canopy structure: Remote sensing of forest disturbance. J. Geophys. Res. 114, (2009).

32. L. A. Rabin, C. M. Greene, Changes to acoustic communication systems in human-altered environments. Journal of Comparative Psychology 116, 137–141 (2002).

33. U. de Camargo, T. Roslin, O. Ovaskainen, Spatio-temporal scaling of biodiversity in acoustic tropical bird communities. Ecography (2019).

34. J. Sueur, Cicada acoustic communication: potential sound partitioning in a multispecies community from Mexico (Hemiptera: Cicadomorpha: Cicadidae). Biological Journal of the Linnean Society 75, 379–394 (2002).

35. A. K. Schmidt, H. Römer, K. Riede, Spectral niche segregation and community organization in a tropical cricket assemblage. Behavioral Ecology 24, 470–480 (2012).

36. C. Q. Stanley, M. H. Walter, M. X. Venkatraman, G. S. Wilkinson, Insect noise avoidance in the dawn chorus of Neotropical birds. Animal Behaviour 112, 255–265 (2016).

37. J. K. Balch, T. J. Massad, P. M. Brando, D. C. Nepstad, L. M. Curran, Effects of high-frequency understorey fires on woody plant regeneration in southeastern Amazonian forests. Philosophical Transactions of the Royal Society of London B: Biological Sciences 368, 20120157 (2013).

38. R. R. de C. Solar, et al., How pervasive is biotic homogenization in human-modified tropical forest landscapes? Ecology Letters 18, 1108–1118 (2015).

39. J. M. Silveira, et al., A Multi-Taxa Assessment of Biodiversity Change After Single and Recurrent Wildfires in a Brazilian Amazon Forest. Biotropica 48, 170–180 (2016).

40. C. Gray, et al., FORUM: Ecological networks: the missing links in biomonitoring science. J Appl Ecol 51, 1444–1449 (2014).

41. M. Longo, et al., Aboveground biomass variability across intact and degraded forests in the Brazilian Amazon. Global Biogeochem. Cycles, 2016GB005465 (2016).

42. M. M. Mollinari, C. A. Peres, D. P. Edwards, Rapid recovery of thermal environment after selective logging in the Amazon. Agricultural and Forest Meteorology 278, 107637 (2019).

43. Z. Burivalova, Ç. H. Şekercioğlu, L. P. Koh, Thresholds of Logging Intensity to Maintain Tropical Forest Biodiversity. Current Biology 24, 1893–1898 (2014).

44. J. C. Su, D. M. Debinski, M. E. Jakubauskas, K. Kindscher, Beyond Species Richness: Community Similarity as a Measure of Cross-Taxon Congruence for Coarse-Filter Conservation. Conservation Biology 18, 167–173 (2004).

45. H. Hillebrand, et al., Biodiversity change is uncoupled from species richness trends: Consequences for conservation and monitoring. Journal of Applied Ecology 55, 169–184 (2018).

46. C. B. Muller, I. C. T. Adriaanse, R. Belshaw, H. C. J. Godfray, The structure of an aphid– parasitoid community. Journal of Animal Ecology 68, 346–370 (1999).

47. D. J. Watts, S. H. Strogatz, Collective dynamics of ‘small-world’ networks. Nature 393, 440–442 (1998).

48. C. Dormann, How to be a specialist? Quantifying specialisation in pollination networks. Network Biology 1, 1–20 (2011).

49. G. Csardi, T. Nepusz, The igraph software package for complex network research. InterJournal, complex systems 1695, 1–9 (2006).

50. J. Oksanen, et al., Community ecology package. R package version 2 (2013).

