## Supplemental Figures for "Animal soundscapes reveal key markers of Amazon forest degradation from fire and logging"

**This PDF file includes:**

Figures S1 to S4

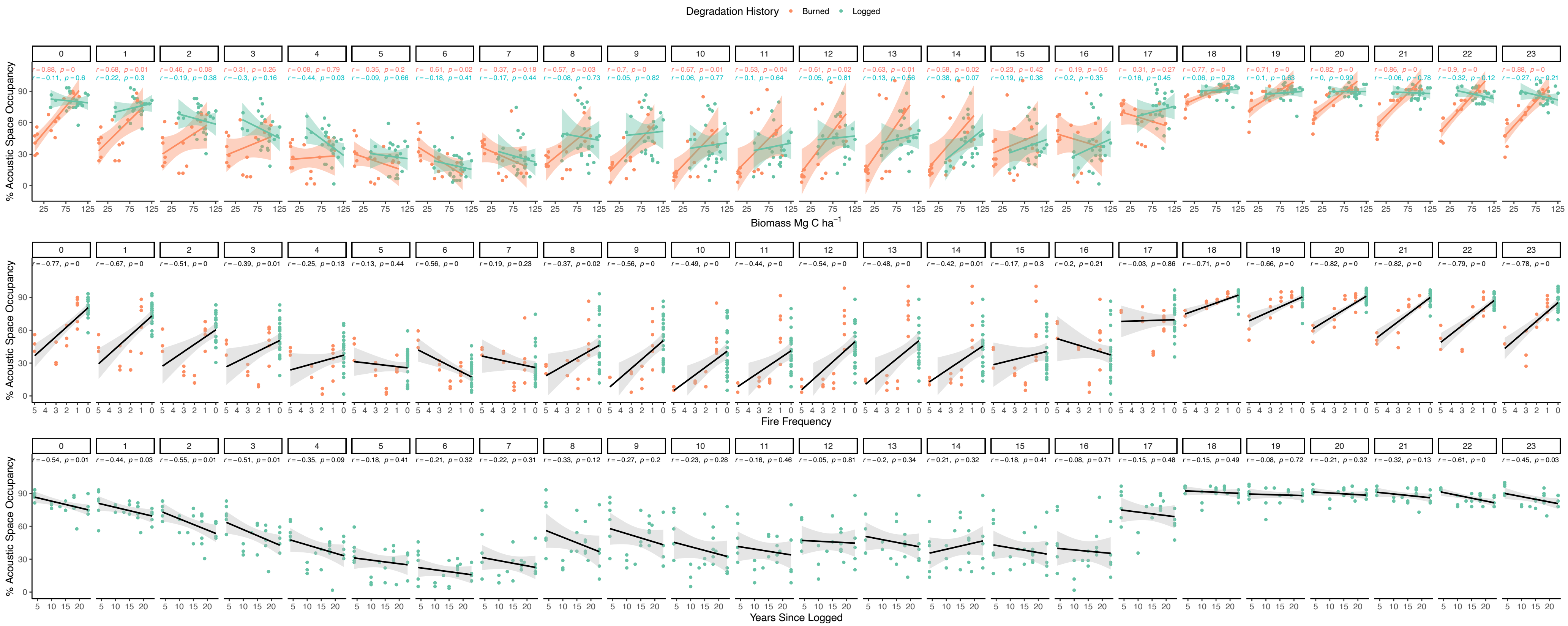


**Figure S1.** Acoustic space occupancy (ASO) aggregated hourly for the 24-hour cycle reveal temporal variation in the dependence of ASO on biomass (top), fire frequency (middle), and years since logging (bottom).

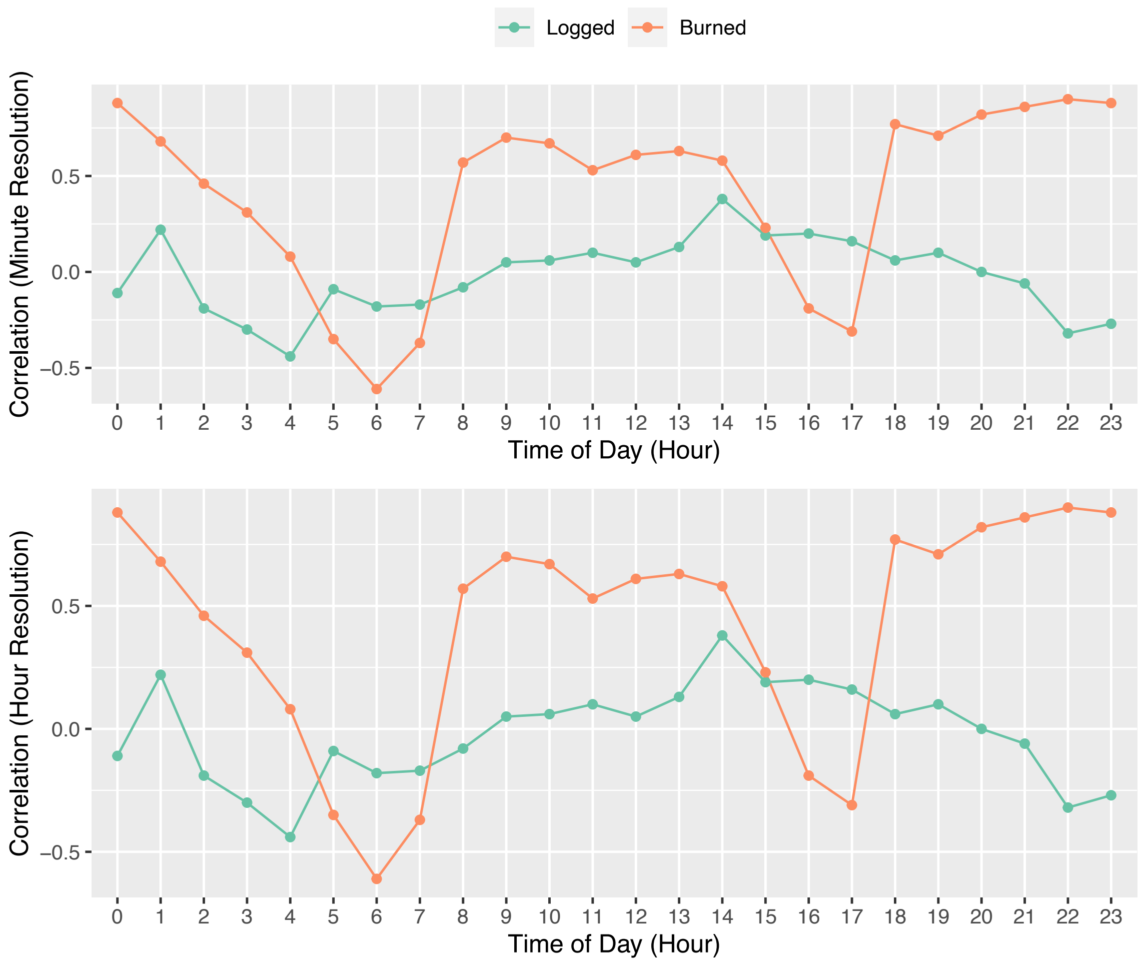


**Figure S2.** Comparison of correlations show robust relationships between degradation and acoustic space occupancy irrespective of the scale of aggregation (hourly vs. minute time steps).

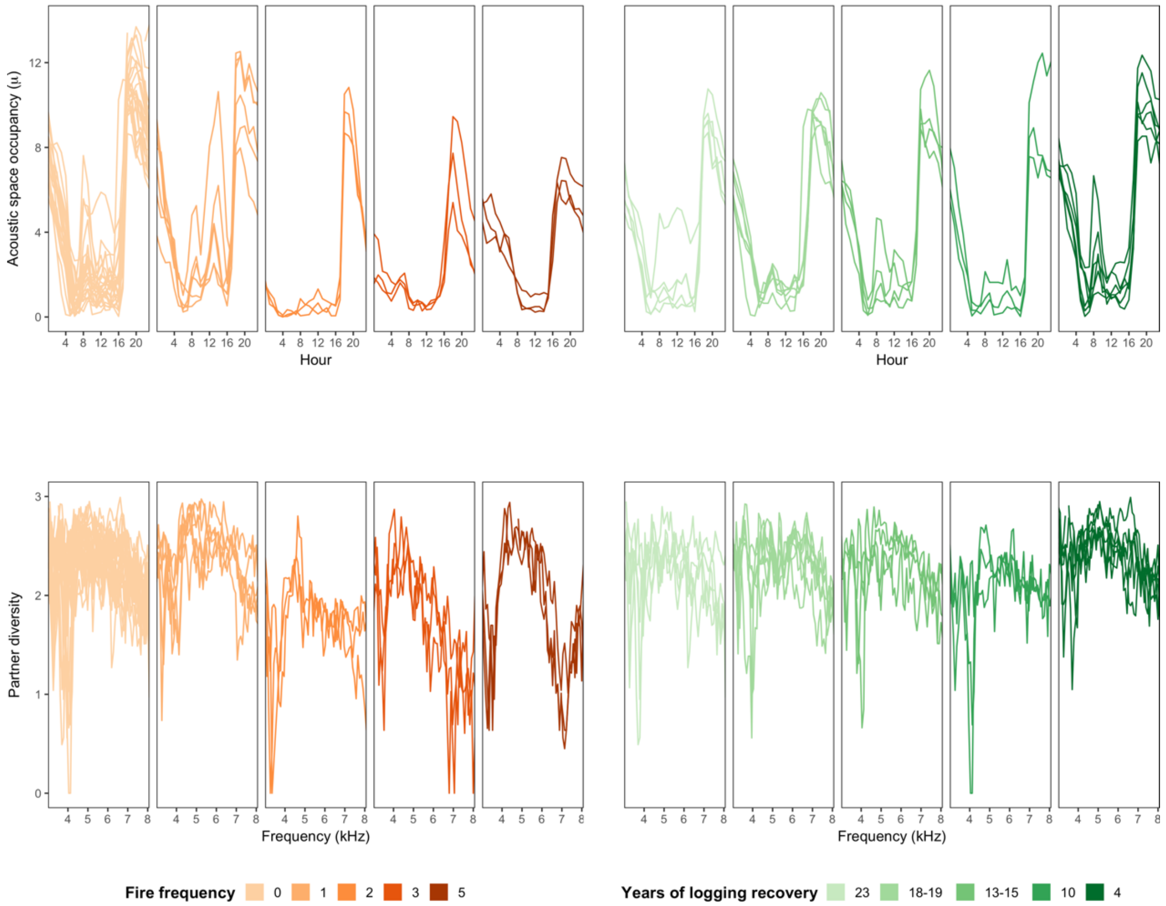


**Figure S3.** Acoustic space occupancy (top) and *partner diversity* (bottom) are highly consistent among site replicates per degradation stratum as a function of fire frequency (left) and post-logging recovery (right).

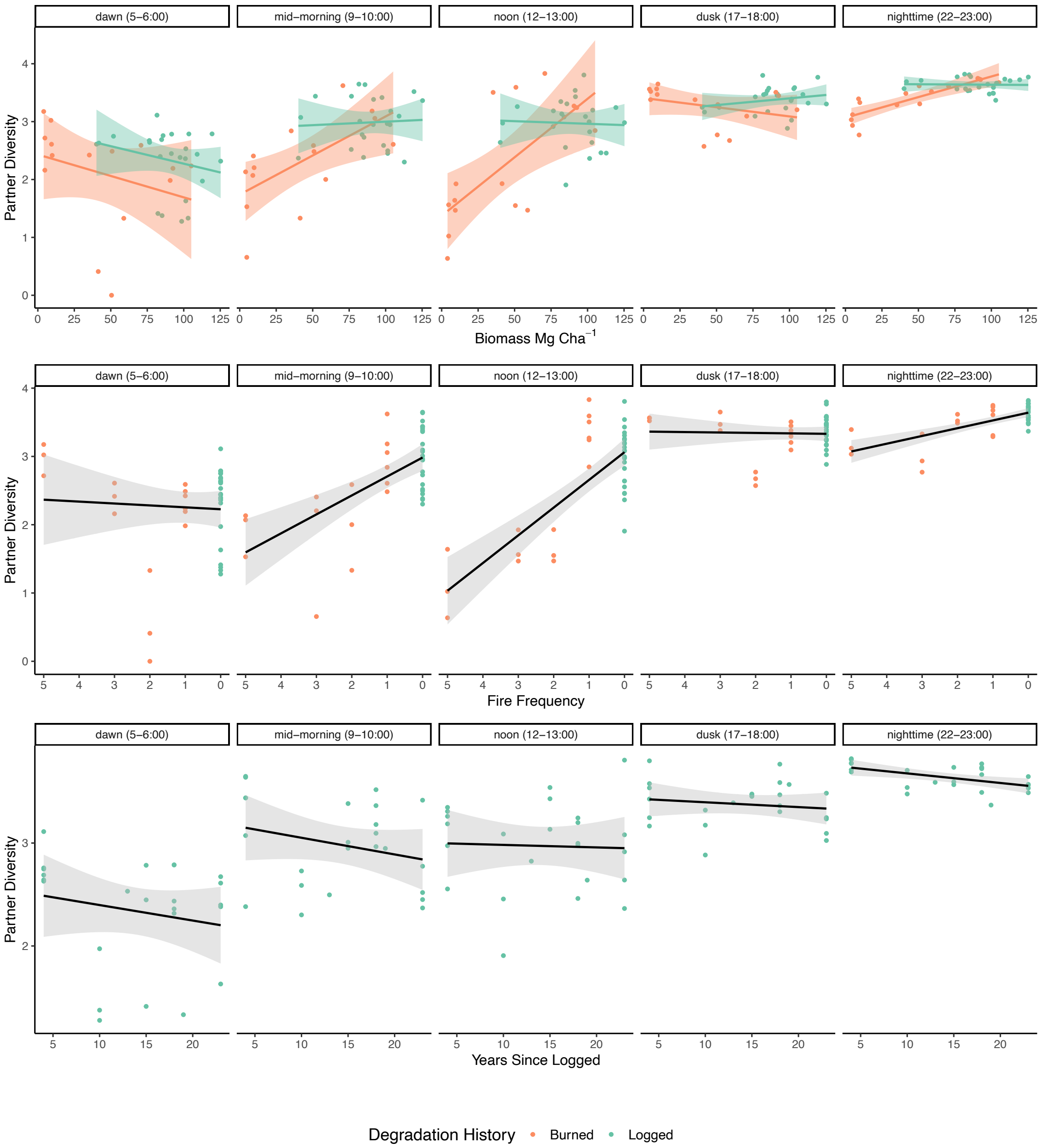


**Figure S4.** Relationships between *partner diversity* of sound hours as a function of biomass (top), fire frequency (middle) and years-since-logging (bottom) depend strongly on time of day. These results are consistent with the ASO-based analysis in Fig. 2.
